# *In planta* exploitation of leaf apoplastic compounds: a window of opportunity for spatiotemporal studies of apoplastic metabolites, hormones and physiology

**DOI:** 10.1101/2023.04.05.535553

**Authors:** Bastian L. Franzisky, Jakob Sölter, Cheng Xue, Klaus Harter, Mark Stahl, Christoph-Martin Geilfus

## Abstract

Processes in the leaf apoplast are relevant for development, cell wall rheological properties, plant nutrition, sink-source portioning, microbe-host plant-interactions or intercellular information exchange and signaling and are therefore regulated or influenced by the composition of the leaf apoplastic solute.

In contrast to the traditional methods for the extraction of apoplastic solutes that are more or less destructive, we propose a new method that allows extraction of leaf apoplastic solutes (i) non-invasively and, thus, (ii) over time. Moreover, the method has (iii) a high spatial resolution that allows identification of solute-microdomains in the leaf apoplast. The method was established for *Arabidopsis thaliana* and *Vicia faba* leaves but should also be applicable to other plants species with similar leaf morphologies. It is based on the infiltration of an aqueous extraction solution into the apoplast followed by its recovery seconds later, both through the stomata. By this, the apoplast (and its solutes) of an identical leaf can be sampled on successive days with negligible symplastic contamination.

A spatiotemporal mapping of leaf apoplastic ion and metabolite patterns within the identical leaf opens a window of opportunity for understanding apoplast biology. As for example, the existence of apoplastic abscisic acid gradients within a leaf in response to salinity was witnessed in this study, as was the unsuspected accumulation of kaempferol glycosides in the leaf apoplast.

The presented method is relevant for plant developmental biologists, phytopathologists, plant physiologists, plant nutritionists and others that need to integrate apoplast biology into their research approaches.

## 1. Introduction

The leaf apoplast comprises the extraprotoplastic matrix including the cell walls, the intercellular gas-filled spaces and the solute-containing (i.e. ions, metabolites, proteins, etc.) fluid film that adheres to the outer surface of the plasma membrane (PM) and surrounds parts of the cell wall fibrils (Dietz, 1997; Ernst Münch, 1930). This apoplastic fluid serves various important functions: Transport of water and solutes such as ions (Sattelmacher et al., 1998), metabolites (O’Leary et al., 2016), proteins (Witzel et al., 2011) or compounds for short- and long-distance signaling. Examples are abscisic acid (Li et al., 2018), electric signals (Zimmermann et al., 2009), reactive oxygen species (Eljebbawi et al., 2021; Podgórska et al., 2017), pH (Geilfus, 2017), and Ca^2+^ (Lionel et al., 1998). Apoplastic signaling is therefore relevant for plant development and response to environmental cues (O’Leary et al., 2016; Rodríguez-Celma et al., 2016; Castro et al., 2021; Geilfus, 2017).

Despite being an compartment relevant for signaling, the apoplast is considered a storage not only for ions [apoplast as polarized ion ‘exchanger’ (Grignon and Sentenac, 1991; Sattelmacher, 2001; Sattelmacher et al., 1998)] but also water (James et al., 2006) and possibly CO_2_ (HCO_3_^-^), i.e. in the sub-stomatal cavity (Santos et al., 2021). Moreover, many specialized processes take place in the apoplast, including the activation of the apoplastic-located class III peroxidases, which play a role in withstanding biotic attacks (Baker et al., 2012; Plieth, 2012), and long-distance movement of miRNAs through apoplastic transmission (Marín-González and Suárez-López, 2012). Last but not least, the apoplast is a refuge for beneficial, parasitic and pathogenic microorganisms such as bacteria (Geilfus et al., 2020; Romero et al., 2019; Rico et al., 2018) and fungi (Hill and Solomon, 2020; Takeda et al., 2009). The chemistry of the apoplastic fluid, i.e. the composition and concentration of compounds dissolved in the apoplastic fluid, is highly variable and influenced by environmental conditions such as light (Mühling et al., 1995; Pitann et al., 2009; Mühling and Läuchli, 2002), water availability (James et al., 2006), stress (Felle, 2001), microbial colonization (O’Leary et al., 2016) and nutrition (Mühling and Läuchli, 1999).

The apoplastic solute composition is under strict control of the symplast. For instance, the activity of the PM H^+^-ATPase can quickly change the apoplastic proton concentration (Geilfus and Mühling, 2012) and thereby energize the PM, which is a prerequisite for differential cell elongation growth in plant tissues (Falhof et al., 2016; Großeholz et al., 2022) and regulating transport across the PM (Huang et al., 2021; Großeholz et al., 2022; Schepper et al., 2013). Spatial apoplastic concentration gradients can result from differential rates of uptake or efflux from the symplast, as demonstrated by proton concentration gradients in leaf microdomains (leaf teeth) (Canny, 1987; Wilson et al., 1991), increased apoplastic sugar and ion concentrations in the vicinity of closing stomata (Jezek and Blatt, 2017) and proton gradients along the axis of the root tip (Großeholz et al., 2022). Environmental cues such as drought, salt, microbial association (Green et al., 2020) and pathogen infection (Gupta et al., 2015; O’Leary et al., 2016) or ionic stress [e.g. cadmium (Wu et al., 2016), sodium and chloride (Shahzad et al., 2013), aluminum (Poschenrieder et al., 2008)] result in dynamic changes in the ion, protein and metabolite composition of the apoplast (Houston et al., 2016).

Apoplastic solute concentrations can be studied minimal invasively using ion–selective microelectrodes (Zimmermann et al., 2010), microelectrode ion flux estimation (Großeholz et al., 2022), ion selective dyes (Mühling and Sattelmacher, 1997; Fricker et al., 1997) or pH-sensitive green fluorescent proteins (Geilfus et al., 2014; Tran et al., 2022; Barbez et al., 2017). These techniques allow elegant measurements over (a limited) time, their disadvantage is that these techniques are in most cases only suitable to detect a single solvent (e.g. one ion or small metabolite) and there are only sensors for a very limited range of solutes available so far.

Therefore, apoplastic solute concentrations have usually been studied using destructive *in situ* approaches, which have limitations due to time-consuming preparation, limited access to the apoplastic compartment or bias resulting from wounding or cutting (Sattelmacher and Horst, 2007). In developing these ‘traditional’ methods, the authors were well aware of these drawbacks and took great care to prevent these obstacles from becoming too significant (Lohaus et al., 2001). Though destructive, it has to be acknowledged that the extraction of apoplastic solutes *via* washing steps, i.e. the extraction of ‘apoplastic wash fluid’ (AWF), enabled breakthroughs in plant physiology and metabolism. These include insights into abiotic stress responses (Shahzad et al., 2013; Poschenrieder et al., 2008; Sagervanshi et al., 2022) plant-pathogen interactions, e.g. in terms of metabolite availability (Yamada et al., 2016; O’Leary et al., 2016), activation of plant defense responses (Wang et al., 2021) or recruitment of antimicrobial substances *via* extracellular vesicles (Rutter and Innes, 2017).

In using the traditional method, the leaves have to be placed in the barrel of a syringe. Larger leaves have to be cut for this purpose, while smaller leaves need to be plucked. Next, by applying a vacuum, the apoplast of cut or plucked leaves is infiltrated with an ‘infiltration solution’ (Chincinska, 2021), e.g. deionized water for capturing free cations or 50 mM BaCl_2_ for exchangeable ions (Mühling and Läuchli, 1999). The mixture of the ‘infiltration solution’ and the apoplastic fluid that harbors the apoplastic solutes is then extracted by gentle centrifugation. The invasiveness of the method, i.e. cutting or plucking the leaves and further mechanical treatment by vacuum infiltration and centrifugation, might cause inaccuracy and contamination by symplastic substances (Nouchi et al., 2012; Lohaus et al., 2001; O’Leary et al., 2014). To have a proxy for the inevitable symplastic contaminations in the AWFs, cytosolic markers such as the concentration of potassium or the activity of primarily symplastic enzymes (glucose-6-phosphat-dehydrogenase or malate dehydrogenase) are quantified. The invasive AWF extraction approach is not only suboptimal because of symplastic contamination, but also because no further measurements can be carried out on the same leaf afterwards. Furthermore, the plant homeostasis is disturbed by cutting off a leaf, which particularly impairs the subsequent study of systemic stress signals or sink-source relationships on other plant parts.

To aid research on leaf apoplast biology, we present a non-destructive method that allows collecting apoplastic solutes minimal-invasively over time. The ‘infiltration solution’ is infiltrated through open stomata into the leaf apoplast where it mixes with the leaf apoplastic fluid without cutting or plucking the leaf. A few seconds later, this mixture is extracted as AWF through the same leaf pores without any centrifugation step. Both the infiltration and extraction are carried out by using a needle-less syringe. The leaf remains intact, being only minimally disturbed by a procedure that lasts only a few seconds. The new approach allows for the first time repeated day-by-day extraction of AWFs from the identical leaf and, of note, from leaf microdomains enabling the analysis of spatiotemporal gradients of apoplastic solutes. This allows capturing of dozens to hundreds apoplastic solutes and the identification of metabolites that so far have received little attention in apoplast biology.

## 2. Methods

### 2.1 Plant material and growth conditions

*Arabidopsis thaliana* L. ecotype Col-0 and *Vicia faba* L. variety Allison (Norddeutsche Pflanzenzucht Hans-Georg Lembke KG, Hohenlieth, Germany) were used as experimental organisms.

#### Vicia faba

Seeds were imbibed in in aerated CaSO_4_ (0.5 mM) solution for one day at room temperature and then placed in moistened quartz sand. After 10 days, the seedlings were transferred into the hydroponic system. Four plants were placed in 5 L plastic pots containing 4.5 L one-fourth strength nutrient solution. In the following days, the strength of the nutrient solution was increased by one-fourth per day until the final concentration was reached. The full-strength nutrient solution had the following composition: 0.1 mM KH_2_PO_4_, 1.0 mM K_2_SO_4_, 2.0 mM Ca(NO_3_)_2_, 0.5 mM MgSO_4_, 0.06 mM Fe-EDTA, 10 µM NaCl, 10 μM H_3_BO_3_, 2.0 μM MnSO_4_, 0.5 μM ZnSO_4_, 0.2 μM CuSO_4_, 0.1 µM CoCl_2_, 0.05 µM (NH_4_)_6_Mo_7_O_24_. The plants were grown in climate cabinets (WEISS HPS1500/S2, Weiss Umwelttechnik, Heuchelheim, Germany) using climate conditions of 14:10 hr day:night; 22°C, about 65% rel. humidity, 200 µmol photons m^-2^ s^-2^ above canopy. After 35 days of cultivation, NaCl was added to the nutrient solution until a final concentration of 30 mM. AWFs were taken from the plant material starting at 36 days of age (Fig. **S1a**).

#### Arabidopsis thaliana

Seeds were germinated in S1 germination soil (Klasmann-Deilmann GmbH, Geeste, Germany) covered with a transparent lid in the greenhouse. After 10 days, the seedlings were transferred into the hydroponics setup essentially as described above. Six plants were placed in 5-liter plastic pot containing 4.5 L quarter-strength of the above nutrient solution. In the following day, the strength of the nutrient solution was increased to half-strength, which was the final concentration for *Arabidopsis* culture. *Arabidopsis* plants were grown in two experimental groups (Fig. **S1b**), (i) with optimal nutrient supply and (ii) with low magnesium addition (low Mg^2+^). The low Mg^2+^ group received no MgSO_4_ for the first two weeks and was then supplied with 0.125 mM MgSO_4_. After 49 days of cultivation, half of the *Arabidopsis* plants with optimal Mg^2+^ supply was exposed to a 50 mM NaCl. AWFs were taken starting from day 50.

### 2.2 In planta spatiotemporal extraction of leaf apoplastic solutes

The following procedure was established on *V. faba* and *Arabidopsis*. The procedure is visualized in the attached video (Video **S1**).

1. A barrel of a 1 mL needleless syringe is filled with 800 µl of an aqueous (double-distilled water) ‘infiltration solution’ that contains 1 mM of the fluorophore pyranine (pKa in aqueous solution: 7.3; Merck, Darmstadt, Germany). The water serves for extracting the apoplastic solutes, while addition of pyranine is important for another reason: the fluorophore is used as a marker to quantify the proportion of water from the ‘infiltration solution’ that is either lost (i.e. evaporated through the stomata or being taken up into the symplast) or came from the plant internal water pool.
2. The plunger is locked on the barrel and air bubbles are squeezed out.
3. The tip of the barrel, of note without needle, is placed perpendicularly on the top of the leaf.
  - Note: Find out which face of your leaf has the stomata/higher stomatal density. Place the tip on this leaf surface.
4. Next, the ‘infiltration solution’ is infiltrated into the leaf apoplast through the opened stomata by applying gentle pressure on the plunger with the thumb. The index finger of the same hand is put on the underside of the leaf below the syringe’s tip. This handling ensures that tip and leaf are close together; otherwise the ‘infiltration solution’ would squirt out.
  - Note: The flooded apoplastic area appears darkish-green (Fig. **1a,c**).
  - Note: To facilitate infiltration, it is important that stomata are open. In the presented work, sampling was carried out four hours after and before turning the lights on and off, respectively. If the fluid cannot be inserted easily, the stomata might be closed and the method should not be used in this case.
5. The ‘infiltration solution’ and the leaf apoplastic fluid with its solutes are now mixed. Removal of this mixture is done about 10 seconds after infiltration. This is done by applying a gentle negative pressure in the barrel by carefully pulling up the plunger with the thumb (see also Video **S1**). The darkish-green discoloration of the apoplastic area (caused by infiltration the ‘infiltration solution’, i.e. the water) disappears. The extract is ready for analysis or can be frozen; make aliquots if necessary.
  - Note: After applying the negative pressure, it can be observed that the mixture flows back into the barrel. About 90% of the originally infiltrated quantity might be extracted.
  - Note: Use different syringes to avoid contamination between different replicates.
6. Repeating steps 1. to 5. at the identical leaf at different days allows the analysis of the leaf apoplast over time.
  - Note: With larger leaves such as those of *V. faba*, it is possible to infiltrate a confined apoplastic area and not the entire apoplast. Partial infiltration can be monitored via the darkish-green appearance of the flooded part. In this way, for example, the leaf tip apoplast and the leaf base apoplast can be examined individually (Fig. **1d**). This allows conclusions about apoplastic microdomains.

**Figure 1.**
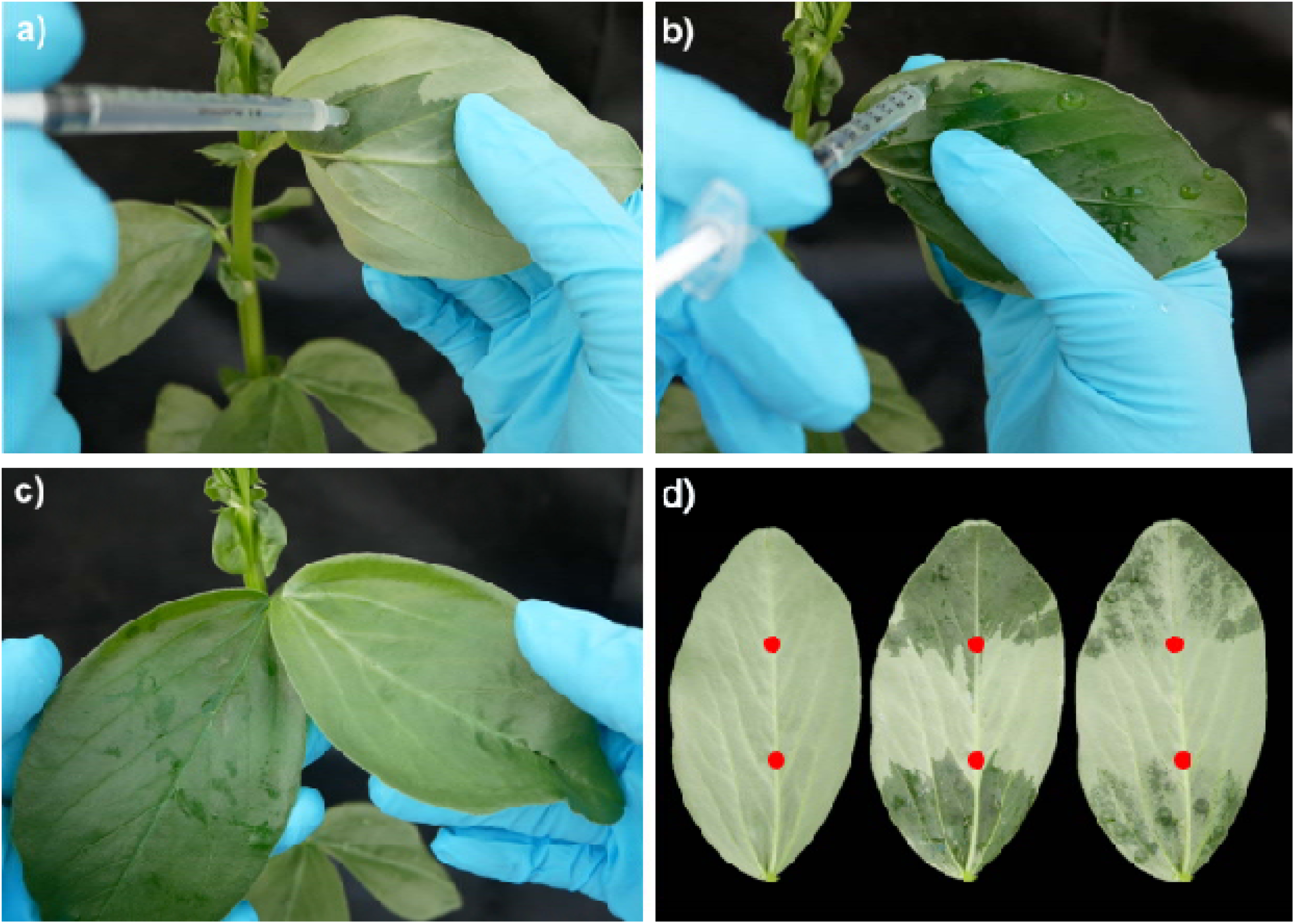
Sampling of apoplastic wash fluid from plant leaves applying the *in planta* extraction method on *Vicia faba*. Infiltration of the ‘infiltration solution’ from the abaxial side of the leave using a syringe (**a**). Infiltrated parts of the leaf can be recognized by the darkish-green color at both abaxial (**b**) and adaxial (**c**) face of the leaf. Infiltration can be confined to specific areas of the leaf (**d**; i.e. partial infiltration for analyzing apoplastic microdomains: here leaf tip *versus* leaf base). Left leaf, unloaded; middle leaf, shortly after loading; right leaf, immediately after extraction of the infiltration solution. Labels (red dots on leaf vein) can be used to tag the relevant position for the following sampling days. This is practical, otherwise the position might be lost.

A challenge that needs to be tackled is that it is never known how much of the ‘infiltration solution’ evaporates through the stomata or is taken up in the symplast. This would lead to overestimation of the solute concentrations. This might happen to a different extent between experimental groups when the treatments differentially influence hydraulic or osmotic parameters in the cell (e.g. salt treatment *versus* control). To account for this, the ‘infiltration solution’ was spiked with 1 mM pyranine (Merck). The pyranine absorption of each AWF was measured to normalize for potential dilution or concentration effects (see 2.5).

A problem with both the traditional ‘infiltration-centrifugation’ method (Lohaus, et al., 2001) and the method described here is that the extracted sample volume is quite low. Therefore, pooling of the samples is recommended. For *Arabidopsis*, AWFs from about 10 to 15 leaves from a total of six plants were combined to obtain a volume of approx. 250 µl, whereas for *V. faba*, AWFs of three leaflets from the same leaf were sufficient.

### 2.3 Assays for checking symplastic contamination

The activity of the symplastic marker enzyme glucose-6-phosphate dehydrogenase (G6PDH, EC 1.1.1.49) in the AWFs was measured according to Borniego et al. 2019. In brief, AWF was centrifuged for 15 min at 15,000 g at 4°C, the supernatant was collected and used for the enzymatic assay. A 10 µl aliquot of the sample was added to 100 µl reaction mix containing 100 mM Tris-HCl pH 8, 6.7 mM MgCl_2_, 12 mM glucose-6-P, 0.4 mM NADP^+^ and incubated at room temperature. The reduction of NADP^+^ to NADPH was followed spectrophotometrically at wavelength of 340 nm. Leaf extracts were generated by homogenization in precooled buffer containing 100 mM Tris-HCl pH 8, 4 mM phenylmethylsulfonyl fluoride; benzylsulfonyl fluoride.

### 2.4 Ion analysis

Concentrations of potassium (K^+^), sodium (Na^+^) and magnesium (Mg^2+^) were measured by using an atomic absorbance spectrometer (3300 series; Thermo Fisher Scientific, Dreieich, Germany), according to the manufactureŕs instructions. Chlorine (Cl^-^) concentration was measured using ion chromatography on a Dionex ICS-2100 (Thermo Scientific) equipped with a Dionex IonPac AS17 column (Thermo Scientific).

### 2.5 Quantification of pyranine as internal standard

Pyranine (8-Hydroxy-pyren-1,3,6-trisufonic acid, also abbreviated as HPTS) was used as an internal standard for normalization of the metabolite results. Sample aliquots of 10 µl were diluted with 200 µl water. Pyranine absorption was determined at 405 nm by using an Epoch microplate spectrophotometer (BioTek/Agilent Technologies, Waldbronn, Germany).

### 2.6 Metabolite analysis

For metabolite analysis, the frozen samples were shipped on dry ice to the Center of Plant Molecular Biology (ZMBP) and stored at – 80 °C until analysis. Amino acid, organic acid, carbohydrate and alcohol concentrations of the AWF samples were quantified by using a Shimadzu TQ 8040 GC/MS instrument. Flavones, amino acids and organic acids were analyzed by using a Waters Acquity-SynaptG2 LC/MS system in addition. Plant hormone analyses were conducted by using a microLC (Trap and Elute M5)–QTrap6500+ (Sciex) system.

For GC/MS analysis 6000 pmol internal standard (3-O-Methyl-d-glucose, OMG) was added to 60 µl AWF sample and dried down with a vacuum concentrator overnight. For derivatization 50 µl Methoxamin (20 mg/ml in Pyridin) was added, ultrasonicated for 10 min and afterwards incubated for 90 min at 30 °C. In a second step 70 µl N-Methyl-N-(trimethylsilyl)-trifluor-acetamid (MSTFA) was added and samples were incubated for 60 min at 40 °C, afterwards kept for another 2 h at room temperature. A volume of 1 µl was analyzed in splitless mode on a Restek SH-Rxi-5SIL-MS column. For separation a He flow rate of 0.92 ml min^-1^ and a temperature gradient of 10 °C min^-1^ from 100 to 320 °C was used. The mass spectrometer was operated in EI ionization and MRM mode.

For untargeted LC/MS analysis, 3 µl of the AWF samples were directly injected onto a Waters Acquity C_18_ SB 150 x 1,0 mm, 1,8 µm column. Separation was performed at 30 °C and a flow rate of 50 µl min^-1^ using a 10 min gradient from 99 % water to 99 % methanol, both solvents containing 0.1 % formic acid. The SynaptG2 mass spectrometer was operated in positive and negative electrospray ionization mode. MS and MS^E^ spectra were recorded in parallel from m/z 50 to 600 using a scan rate of 0.5 seconds and a resolution of 10000. For compound quantification extract ion chromatograms were generated and integrated.

For plant hormone analyses, 25 µl of the AWF samples were diluted with 50 µl chilled 0.1 % formic acid and 25 µl 80 % methanol (containing the isotopically labeled internal standards of jasmonic acid, abscisic acid, indole-3-acetic acid (50 nM each) and salicylic acid (25 nM)). 50 µl of the solution were injected onto a Phenomenex Luna C_18_ trapping column (20 × 0.3 mm, 5 µm, flow rate: 25 µl min^-1^) and afterwards separated on a Phenomenex Luna Omega Polar C_18_ column (150 × 0.3 mm; 3 μm) at 55 °C and a flow rate of 10 µl min^-1^. For this the following gradient was used: 0 - 0.2 min, isocratic 90 % A; 0.2 – 2 min, linear from 90 % A to 30 % A; 2 - 4.5 min, linear from 30 % A to 10 % A; 4.5 – 5 min, linear from 10 % A to 5 % A; 5 - 5.3 min, isocratic 5 % A; 5.3 - 5.5 min, linear from 5 % A to 90 % A; 5.5 – 6 min, isocratic 90 % A (A: water, 0.1 % formic acid; B: acetonitrile, 0.1 % formic acid). The QTrap 6500+ mass spectrometer was operated in positive and negative electrospray ionization mode (Optiflow Turbo V ion source) using the MRM transitions given in Table **S1a**.

### 2.7 Experimental setup and data analysis

To validate the robustness of the new AWF extraction method, various experiments were conducted as shown in Figure **S1**. The AWFs were sampled in randomized order, yielding 6 to 10 biological replicates per variable (Table **S1b,c**). To account for the leaf-specific dilution effect (Lohaus et al., 2001), all AWF-derived data, i.e. ion concentrations, metabolite peak areas and enzyme activities, were normalized to the pyranine absorption intensity of each single AWF before further analysis. Data were analyzed using R (R Development Core 2017). Individual plants were repeatedly sampled on consecutive days. Due to this repeated-measures design, data was analyzed using the repeated measures algorithm of “lme4” (Bates 2015). Where applicable, ‘treatment’ and ‘days of treatment’ with an interaction term were entered as fixed effects into the model, whereas the ‘plant ID’ was given as random effect to account for the dependency in the data. Missing values in enzyme activity data due to falling below the detection limit of the enzyme assay were replaced with the half of the minimum. The response of phytohormone levels to NaCl treatment was calculated by normalizing to the mean of controls. Residuals of statistical models were inspected visually. Data were analyzed on the basis of *P* ≤ .05 and by using the Tukey test algorithm unless stated otherwise. Data were visualized using ‘ggplot2’ (Wickham 2016).

## 3. Results

### 3.1 Non-destructively sampled AWFs show minimal symplastic contamination

In order to verify that the improved AWF extraction method is minimal-invasive with regard to cell integrity, symplastic markers were monitored such as K^+^ concentration and G6PDH-activity (Baker et al., 2012; Boudart et al., 2005; O’Leary et al., 2014; Lohaus et al., 2001). The detection of larger amounts of these markers would indicate that the method compromises membrane integrity, leading to the leakage of symplastic components into the apoplast.

In the cytosol of plant cells, K^+^ concentration ranges between 100 and 150 mM (White and Karley, 2010). In the apoplast, K^+^ concentration is significantly lower being about 10 mM [e.g. 3 to 12 mM in *V. faba* (Lohaus et al., 2001; Sagervanshi et al., 2022); to our knowledge, no data available for *Arabidopsis*]. Figure **2** shows information about both symplastic marker, K^+^ concentration and G6PDH-activity, in leaf AWFs that were collected from control- and salt-treated *V. faba* and *Arabidopsis* leaves. AWFs extracted from non-stressed control plants showed very low K^+^ concentration (< 1 mM for *V. faba,* Fig. **2b,d**; < 4 mM for *Arabidopsis,* Fig. **2f**), even when plants were exposed to NaCl stress for one day, K^+^ concentration in AWFs of both species did not increase.

**Figure 2.**
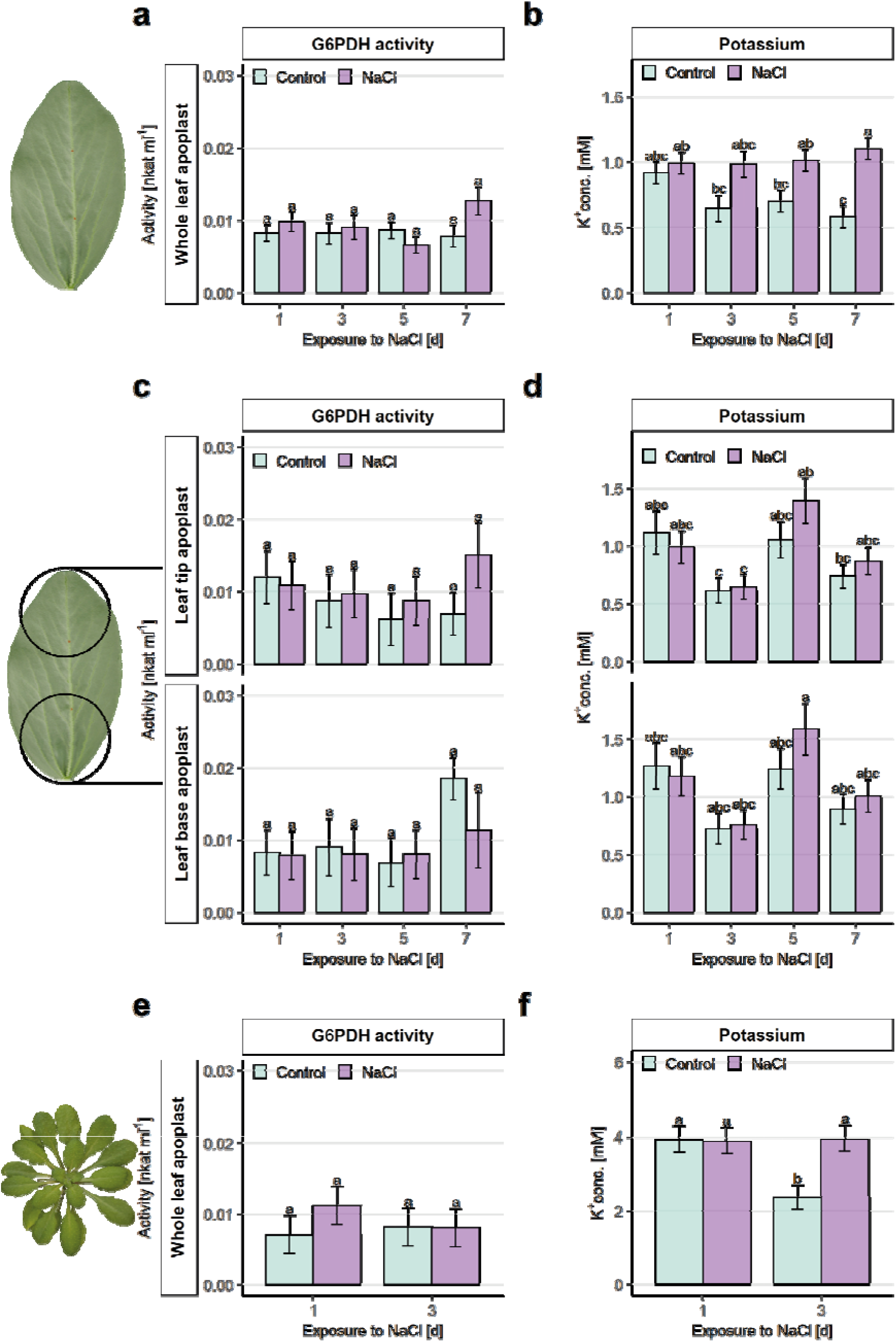
Apoplastic wash fluids (AWFs) show minimal symplastic contamination after repeated sampling of the identical leaf, irrespective of a mild salt treatment. The markers for symplastic contamination, i.e. the activity of the enzyme glucose-6-phosohate dehydrogenase, and the K^+^ concentration were measured in AWFs from whole leaf apoplast (**a-b**), apoplastic micro-domains of *Vicia faba* leaves exposed to mild salt stress ( 30 mM NaCl) from 35 days after germination (**c-d**), and whole leaf apoplast of *Arabidopsis thaliana* exposed to mild salt stress (50 mM NaCl) from 49 days after germination (**e-f**). The AWFs were repeatedly sampled from the identical leaves over the experimental period indicated as days of exposure to NaCl. The first sampling was done at one day after NaCl exposure, while the repeated extraction of AWFs was done after three days of NaCl exposure for *A. thaliana*, and after three, five and seven days for *V. faba* (Fig. **S1**). Means ± SE (n = 6-8); different letters indicate significant differences between treatments and days of exposure; comparisons within microdomains where applicable (Tukey test*; P* ≤ .05).

Similarly, the G6PDH activity was low in the AWFs extracted from non-stressed controls of both species (< 0.01 nkat g^-1^ FW for *V. faba* and *Arabidopsis*, Fig. **2a,c,e**). Stressing the plants for one day with NaCl did not increase the level of symplastic G6PDH activity (Fig. **2a,c,e**). In addition to the enzyme activity in AWFs, relative activity is often reported, which refers to the activity measured in the leaf homogenate. G6PDH activity of whole leaf homogenates of both species was about 6.5 nkat g^-1^ FW (Table **S1b,c**), corroborating previous reports on symplastic G6PDH activity of 7.6 nkat g^-1^ FW in *Arabidopsis* (Haslam et al., 2003) and 6.9 nkat g^-1^ FW in *V. faba* (Turcsányi et al., 2000). In relation to the activities measured in leaf homogenate, the AWFs of both species derived from the first round of extraction had relative activities of less than 0.15 %. These data clearly demonstrate the absence of significant symplastic contaminations in the AWFs from both plant species (White and Karley, 2010).

After three days of NaCl exposure, and thus two days after the first extraction, AWFs were collected again from the identical leaves of both species. However, after this second round of AWF collection, the leaves of *Arabidopsis* partially showed slight marks from the syringe tip. For this reason, AWF sampling from *Arabidopsis* was not performed a third time. This, however, was doable for *V. faba*: The identical leaf could easily be sampled four times at two-day intervals. Both symplastic markers were also analyzed in the AWFs after repeated sampling, in both the NaCl-stressed and the non-stressed control plants. The repeated sampling from identical leaves did not cause a significant increase in K^+^ concentrations (Fig. **2b,d,f**) and G6PDH activity (Fig. **2a,c,e**) in the AWFs of both species. In contrast, there was a trend to a decrease in the K^+^ concentration in the AWFs of non-stressed controls (Fig. **2b**) suggesting that the repeated extractions deplete the apoplast for solutes such as K^+^ (see section 4.2). Such a ‘washout effect’ was far less observed for the G6PDH activity (Fig. **2a**), probably because the apoplastic presence of the enzyme is negligible (i.e. nothing can be washed out). However, this is different for apoplastic K^+^ whose loss probably cannot be fully compensated by replenishing ions from the symplast. Importantly, to account for bias due to a putative washout-effect and to allow straightforward inter-species comparison of apoplast responses, the concentrations of all the apoplastic compounds presented next (ions, metabolites, and phytohormones) were normalized. For this purpose, the values measured in the NaCl treatment were divided by their controls on each sampling day (expressed as log_2_ fold-changes). This procedure is recommended for users who aim to repeatedly extract AWF from identical leaves, *viz*. for users that might face such a ‘washout effect’.

### 3.2 Concentration of Na^+^, Cl^-^ and Mg^2+^ in the leaf apoplast depends on the availability of these ions at the roots

To demonstrate that the new protocol is suitable for depicting plant nutritional processes in the leaf apoplast, we analyzed the ion composition in AWFs prepared from *Arabidopsis* plants grown under optimal magnesium (0.5 mM Mg^2+^) or sub-optimal magnesium (0.125 mM Mg^2+^) nutrition. The *Arabidopsis* plants that were optimally supplied with Mg^2+^ showed concentrations of about 1 mM Mg^2+^ in the AWF (Fig. **3**), while sub-optimal Mg^2+^ supply resulted in a halved Mg^2+^ concentration (0.5 mM). These AWFs had low levels of symplastic contamination as well (Table **S1d**).

**Figure 3.**
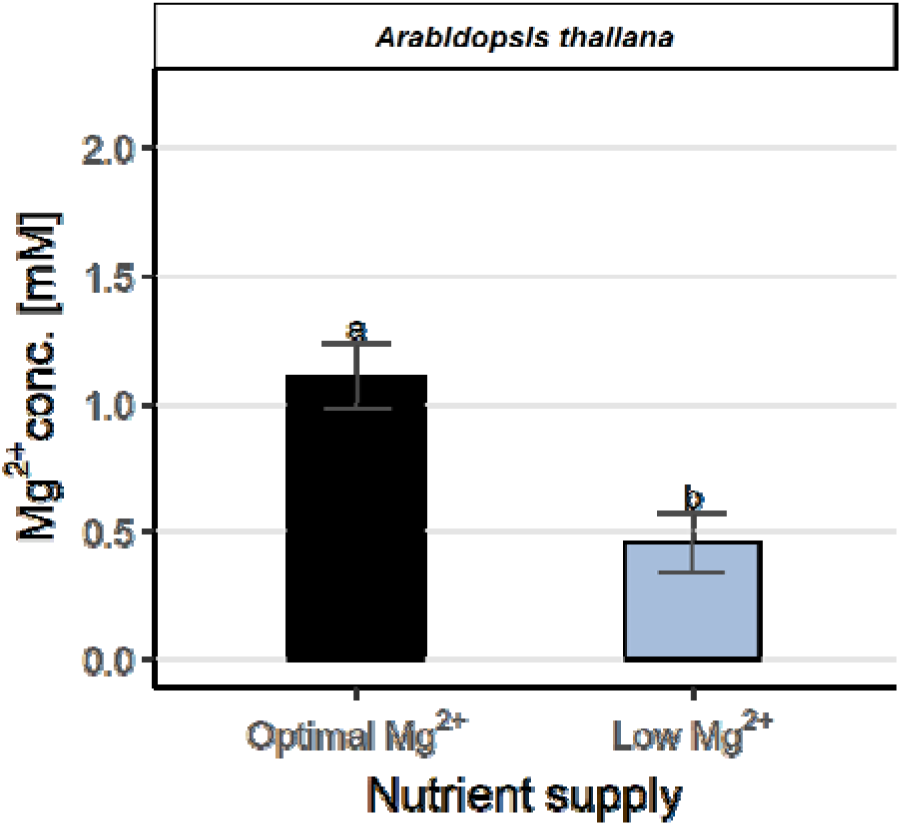
Low Mg^2+^ supply at the roots leads to lower Mg^2+^ concentrations in the leaf apoplast. Mg^2+^ concentrations in AWFs from whole leaf apoplast of *Arabidopsis thaliana. A. thaliana* was grown with optimal (0.5 mM Mg^2+^) or sub-optimal (0.125 mM Mg^2+^) Mg^2+^. AWFs were extracted 50 days after germination (Fig. **S1**). Means ± SE (n = 5); different letters indicate significant differences between comparisons of Mg^2+^ supply variants (Tukey test*; P* ≤ .05).

The salt stress experiment (Fig. **S1**) can serve as a proof of concept to monitor ― in this case ― stress-physiological processes over time. The experiment revealed that when NaCl was applied to the plants, the concentrations of Na^+^ and Cl^-^ increased in the leaf AWFs of both species (Fig. **4**). By repeatedly extracting AWFs from identical leaves, it was found that the concentration of Na^+^ in the whole leaf apoplast of *V. faba* increased by 0.3-fold two days after the onset of stress and by 0.5-fold after seven days (Fig. **4a**), while Cl^-^ increased by up to 0.6-fold after seven days (Fig. **4b**). Similarly, in *Arabidopsis*, the Na^+^ and Cl^-^ concentrations increased by 2-fold and 2.2-fold, respectively, after three days stress treatment (Fig. **4c,d**). *Arabidopsis* accumulated more Na^+^ and Cl^-^ ions in the leaf apoplast than *V. faba*, which is in agreement with the different NaCl doses applied to each species, i.e. 30 mM NaCl for *V. faba* and 50 mM NaCl for *Arabidopsis*. These results show that the new method is suitable for monitoring ion accumulation in the leaf apoplast over extended periods by repeated sampling of identical leaves.

**Figure 4.**
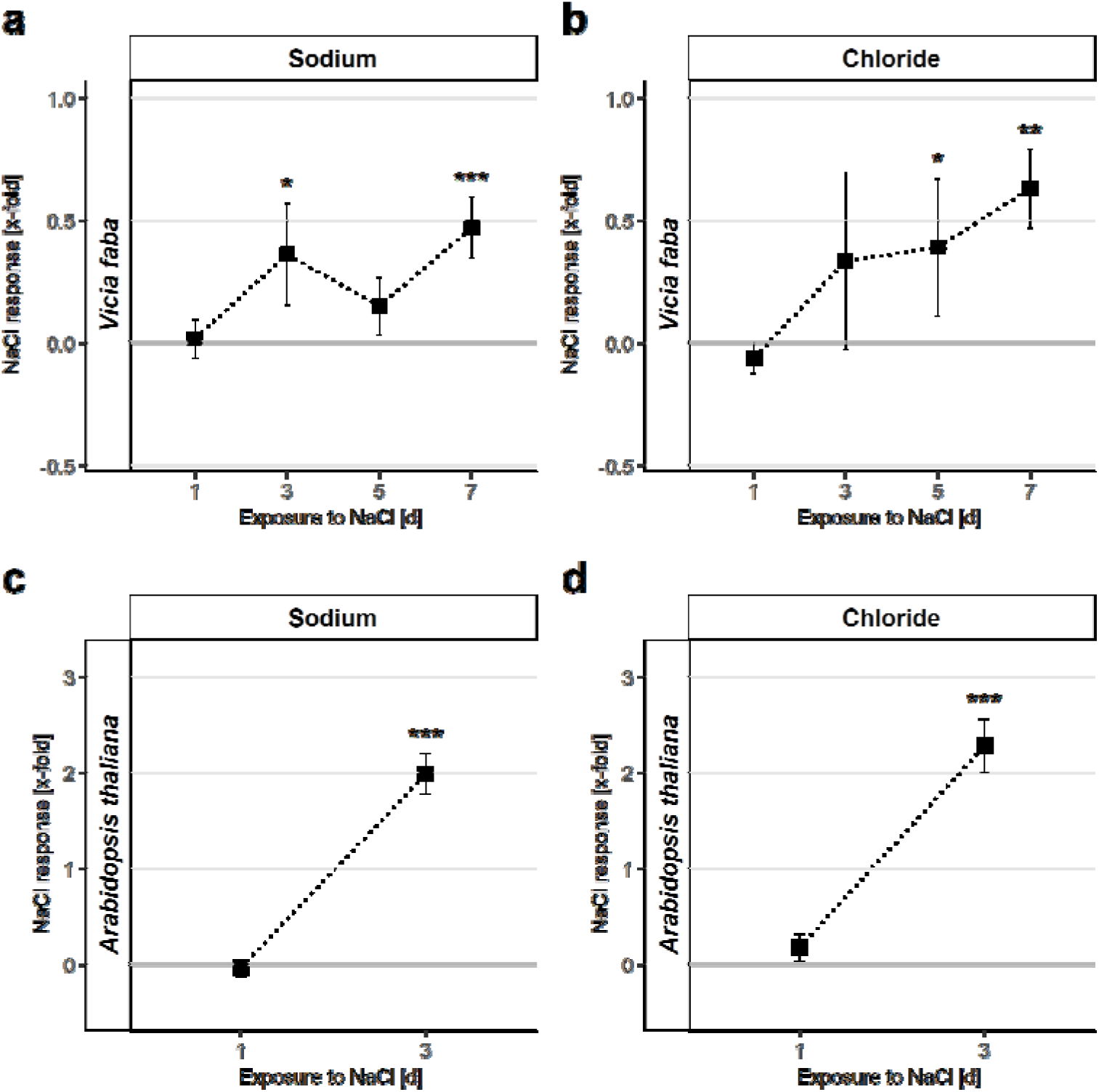
Na^+^ and Cl^-^ accumulate in the apoplast of *Vicia faba* and *Arabidopsis thaliana* after exposure to mild NaCl stress. Changes in concentrations of Na^+^ and Cl^-^ in whole leaf AWFs from *V. faba* (**a-b**) and *A. thaliana* (**c-d**) after exposure to NaCl treatment for 7 days (*V. faba*) or 3 days (*A. thaliana*). The AWFs were repeatedly sampled from the identical leaves over the experimental period indicated as days of exposure to NaCl. The first sampling was done at one day after NaCl exposure, while the repeated extraction of AWFs was done after three days of NaCl exposure for *A. thaliana*, and after three, five and seven days for *V. faba. V. faba* was exposed to 30 mM NaCl at 35 days after germination, while *A. thaliana* was exposed to 50 mM NaCl at 49 days after germination; controls without salt stress as shown in Fig. S1. Data were normalized over the non-NaCl-stressed control plants, i.e. means of the fold-change_log2(NaCl/Control)_ ± SE (n = 6-9). Asterisks indicate the level of significance of comparisons between NaCl treatment and control (*, *P* ≤ .05; **, *P* ≤ .01; ***, *P* ≤ .001).

### 3.3 The presented protocol enables to monitor apoplastic metabolite dynamics in the identical leaf over time

Applying any of the traditional established methods, it is not possible to analyze the dynamics of solutes in the apoplast of an identical leaf over time. In order to show that this is possible with the new method presented here, the temporal course of apoplastic metabolites was monitored in response to salinity in *V. faba* and *Arabidopsis*. Metabolite detection was done by using liquid chromatography mass-spectrometry (LC/MS) and gas chromatography mass-spectrometry (GC/MS). The combination of these approaches enabled the identification and quantification of several metabolite groups in the AWFs of both species: sugars (e.g. fructose, raffinose, sucrose, etc.), various amino acids (e.g. ɣ-amino butyric acid (GABA), beta alanine, glutamine, leucine, lysin, phenylalanine, serine, etc.), polyols (glycerol, mannitol, and sorbitol) and two flavones, most likely the bisglycosides kaempferithrin and kaempferol glucoside rhamnoside (Fig. **5**). Moreover, organic acids (citric-, fumaric-, malic-, succinic acid, etc.) and several hormones such as indole-3-acetic-, salicylic-, jasmonic- and abscisic acid were identified in the AWFs of both species.

**Figure 5.**
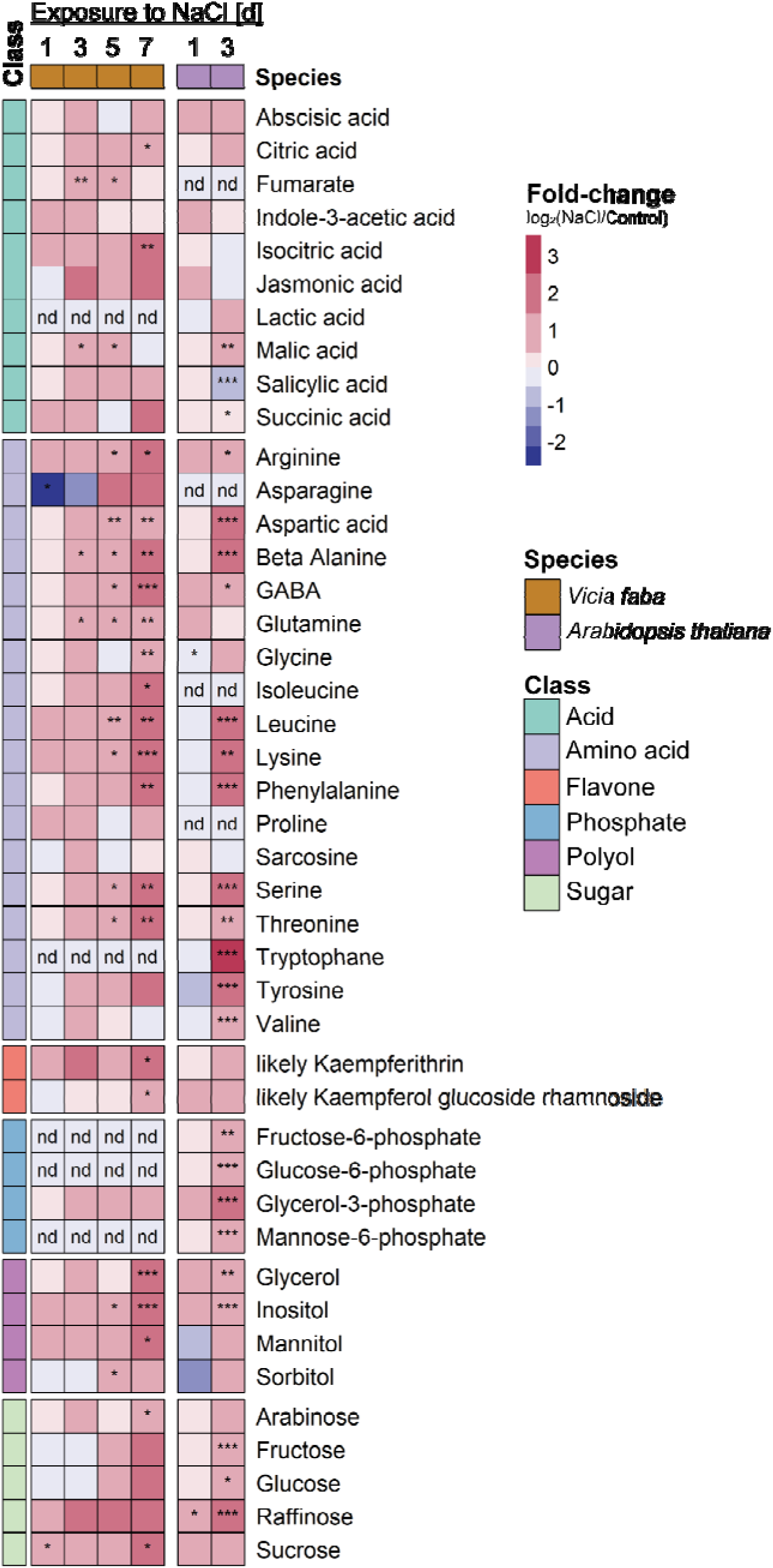
Metabolic response to mild salt stress in the leaf apoplast of *Vicia faba* and *Arabidopsis thaliana*. AWFs taken from whole leaves of *V. faba* and *A. thaliana*. The AWFs were repeatedly sampled from the identical leaves over the experimental period indicated as days of exposure to NaCl. The first sampling was done at one day after NaCl exposure, while the repeated extraction of AWFs was done after three days of NaCl exposure for *A. thaliana*, and after three, five and seven days for *V. faba. V. faba* was exposed to 30 mM NaCl at 35 days after germination, while *A. thaliana* was exposed to 50 mM NaCl at 49 days after germination as shown in Fig. **S1**. Data were normalized over non-NaCl-stressed control plants. Mean of the fold-change_log2(NaCl/Control)_ of metabolite contents is indicated by a color code (n = 6-8). Asterisks indicate the level of significance (*, *P* ≤ .05; **, *P* ≤ .01; ***, *P* ≤ .001; GABA, γ-aminobutyric acid; nd, no data available).

In the apoplast of *V. faba* leaves, the metabolic response to NaCl was dominated by an increase of various free amino acids (Fig. **5**). Among them was GABA, which plays a significant role in salt stress acclimatization (Su et al., 2019; Che-Othman et al., 2020; Munns et al., 2020) and regulation of stomata aperture under drought (Xu et al., 2021). Also sugars, such as sucrose and arabinose, and the polyols glycerol and mannitol accumulated apoplastically in leaves of *V. faba* in response to the NaCl treatment. AWFs from *Arabidopsis* leaves showed a similar metabolic response as *V. faba* leaves, except for the increase in hexose phosphates fructose-6-phosphate and glucose-6-phosphate, which was specific for *Arabidopsis*. The NaCl treatment caused accumulation of amino acids and sugars. In line with the higher NaCl dose of 50 mM for *Arabidopsis* (*versus* 30 mM for *V. faba*), these changes emerged faster in *Arabidopsis* than in *V. faba*, i.e., after already three days of salt treatment.

### 3.4 Hormonal microdomains are present in the leaf apoplast

As shown above, the new method is suitable to track leaf apoplastic ion- and metabolite-dynamics in identical leaves over time. Because the sampled leaves remain intact, it was tested next if the apoplast of the tip and that of the base of the same leaf can be analyzed separately. To check this, the normalized concentrations of selected phytohormones such as abscisic acid (ABA), indole-3-acetic acid (IAA), jasmonic acid (JA), and salicylic acid (SA) were monitored after exposure of *V. faba* plants to salt stress over time (Fig. **6**). At the first day of NaCl exposure, ABA increased only in the AWF of the leaf base, while this response was not yet apparent in the AWF of the leaf tip. In contrast, ABA levels in the AWF of the ‘whole leaf did not show this difference, witnessing the need for spatially resolved extraction. Another instance that highlights the utility of a spatially resolved analysis emerged from the IAA determination. When challenged by salinity, the apoplast of the leaf tip responded differently compared to the apoplast of the base or whole leaf. In the AWF of the leaf tip, IAA was reduced 1-fold after one day of salt stress, while it remained unchanged in the AWFs of the leaf base and the whole leaf. In the further course of the salt exposure from day three to seven, there were similar IAA levels in all apoplast fractions. The levels of JA in the AWFs obtained from both the leaf tip and the leaf base were decreased by 1.5-fold after one-day exposure to NaCl. However, after three days of exposure, there was only a 1-fold reduction in the AWFs of both apoplastic sub-compartments, indicating a slight increase in JA levels compared to the previous measurement. At the same time points, JA levels in the AWFs of whole leaf were increased in response to salt. Thus, a contrasting response was visible again. With respect to SA, a similar pattern to JA was observed, with more reduced SA levels in the apoplast microdomains (about −1.5-fold) compared to that of the whole leaf (unchanged). In the following days of NaCl exposure, SA levels reverted to those of the non-stressed controls, except for a slight trend toward an increased SA level in the leaf base apoplast after five days of exposure. Overall, results demonstrate that analyses restricted to the entire leaf might not give a realistic picture of the differential apoplastic conditions within one and the same leaf.

**Figure 6.**
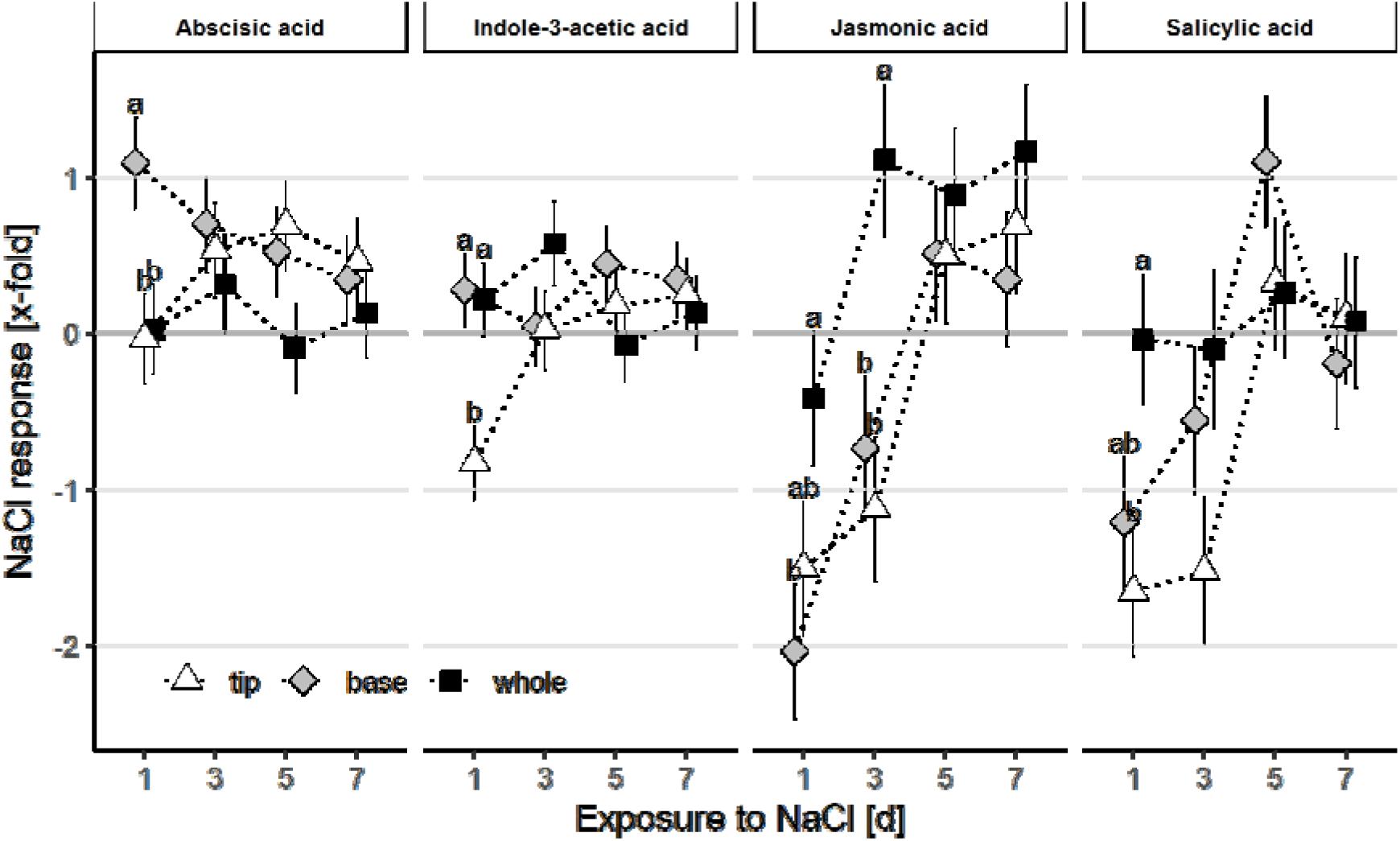
Mapping spatial concentrations of phytohormones concentrations in the leaf apoplast of *Vicia faba* in response to mild NaCl stress. Levels of the phytohormones abscisic acid, indole-3-acetic acid, jasmonic acid, and salicylic acid in AWFs sampled from whole leaves and leave micro domains, i.e. leaf base and tip, of *V. faba* exposed to 30 mM NaCl stress and corresponding non-stressed controls. The AWFs were repeatedly sampled from the identical leaves over the experimental period indicated as days of exposure to NaCl. The first sampling was done at one day after NaCl exposure, while the repeated extraction of AWFs was done after three, five and seven days. Plants were exposed to a 7-day NaCl stress-treatment at 35 days after germination as shown in Fig. **S1**. Data were normalized over mean of non-NaCl-stressed control plants, i.e. means of the fold-change_log2(NaCl/Control mean)_ ± SE (n = 6-9). Different letters indicate significant differences of comparisons between apoplast microdomains within a sampling day (Tukey test*; P* ≤ .05).

## 4. Discussion

### 4.1 A new opportunity for the study of the leaf apoplast arises through the non-invasive collection of leaf apoplastic solutes: acquisition of time series with minimal symplastic contamination

It is increasingly recognized that apoplastic processes are relevant for plant development, morphology, nutrition, physiology and ecology. Research on apoplast biology has increased over the past three decades (Table S1d), demonstrating that there is vibrant plant science community that will benefit from our new method. So far, the main problem concerning the study of leaf apoplast biology was gaining access minimal-invasively to this compartment. Except of sensors that can only measure one variable over a short period of time (hours), e.g. protons (Gjetting et al., 2012; Moreau et al., 2022) or calcium (Gao et al., 2004), the combined simultaneous analysis of several apoplastic compounds from the same leaf over time has not been possible so far. Our new method for the in planta extraction of apoplastic solutes overcomes these limitations: Even the analysis of leaf apoplastic microdomains is doable in the identical leaf over time.

So far, measurements using the traditional ‘infiltration-centrifugation’ technique have been hampered by various issues, including: (i) the inability to sample the same leaf over a period of time, (ii) a lengthy processing time of approximately 5-30 minutes, (iii) mechanical stress caused by the need to centrifuge the leaf and potentially pick or cut longer leaves, and (iv) contamination from the symplast resulting from leaf cutting. Having as less symplastic contaminations as possible is essential for the interpretation of the results. Contamination markers such as K^+^ (Lohaus et al., 2001) and the activity of symplastic enzymes such as malate dehydrogenase (MDH) and glucose-6-phosphate dehydrogenase (G6PDH) are frequently used (Baker et al., 2012; Boudart et al., 2005; O’Leary et al., 2014). AWFs that were collected from non-stressed plant leaves using the traditional ‘infiltration-centrifugation’ technique showed K^+^ concentrations between about 3 to 12 mM K^+^ in *V. faba* (Lohaus et al., 2001; Shahzad et al., 2013; Sagervanshi et al., 2022), pea, spinach (Speer and Kaiser, 1991), and sunflower (Nikolic and Römheld, 2003), while up to about 19 mM K^+^ was reported for common bean (O’Leary et al., 2016). In comparison, the AWFs gained from *V. faba* using the new method had at least 65% lower K^+^ concentration compared to the traditional method (i.e. 1 mM *versus* 3 to 12 mM). This was also true when identical leaves were repeatedly sampled and mild salt stress was applied.

A similar picture emerges when the activity of G6PDH as a contamination marker is examined (Fig. **2**; Table **S1e**): Previous studies using the traditional ‘infiltration-centrifugation’ technique to extract AWFs from non-stressed plants reported G6PDH activities normalized to that of leaf homogenate between 0.1 to 3% for *Arabidopsis* (Rossi et al., 2021; Haslam et al., 2003) and up to 7.2% for *V. faba* (Turcsányi et al., 2000). Therefore, the normalized activities of about 0.15% achieved using the new method also demonstrate very low symplastic contamination. The consistently low contamination confirms that the method is minimal-invasive and thus AWFs can be obtained with high purity, even when AWFs are repeatedly sampled from identical leaves of mildly salt-stressed *V. faba* and *Arabidopsis*.

The aim of this study is to demonstrate that the new method has decisive advantages over traditional methods for AWFs collection. The rational for not directly comparing the traditional *versus* the new method is based on two considerations: First, only single extractions are possible using the traditional destructive method. So no measurements over time on the identical leaf are possible. A comparison between the different extraction methods would only be possible with AWFs extracted at a single time point. Second, the decisive quality criterion of an extraction method is the AWF purity, i.e., low symplastic contamination, which is a proof for the methodological robustness. Therefore, previously reported contamination data obtained with traditional destructive AWF extraction methods are valid references to validate the new AWF extraction method, e.g. our own studies on *V. faba* (Shahzad et al., 2013; Sagervanshi et al., 2022) and common bean (O’Leary et al., 2016).

### 4.2 Treatment-specific recovery rates of the ‘infiltration solution’ and a solute depletion in apoplastic extracts due to ‘washout effect’

Bias with regard to varying solute concentration in the AWFs arising from treatment-specific recovery rates of the ‘infiltration solution’ needs to be considered. This is important because many studies are evaluated in a way that the concentration of the solute of interest in the AWF is compared between the treatment groups. The treatment itself, however, could affect the recovery rate of the infiltrated solution. For instance, a treatment that changes osmotic potential of the cell will influence the amount of water from the ‘infiltration solution’ that flows into the cell (Arndt et al., 2015) and, thus, cannot be recovered. To take this into account, pyranine was added as inert marker (Chincinska, 2021) to the infiltration solution, so that the information about the quantity of pyranine in the AWF could be used for normalization. Another bias to be considered arises from a possible depletion of apoplastic solutes by repeated sampling (’washout effect’), as observed for K^+^ (Fig. **2b**). As a result of four samplings on the identical leaves, a decrease in the apoplastic K^+^ concentration of *V. faba* occurred in the control. It is very likely that such depletion effect will also occur in the treated group as function of repeated AWF extractions; here the plants treated with salt stress. To account for this issue, NaCl-treatment was normalized to control to obtain the treatment-specific responses independent of the inevitable ‘washout effect’. In this respect, the experimenter can decide what is more important: the possibility to acquire time series (new method with repeated extraction from identical leaves) or no washout effect (new method, but single extraction only).

### 4.3 Proof of concept: Monitoring changes in nutrient and metabolite pattern in the leaf apoplast in response to salt stress or magnesium deficiency

To validate the new method, we checked whether typical consequences of a decrease or increase of mineral concentrations in the rooting medium can be monitored in the leaf apoplast. Firstly, a decrease in foliar Mg^2+^ concentration is to be expected under Mg^2+^ starvation (Hermans and Verbruggen, 2005). Accordingly, leaf apoplastic Mg^2+^ concentrations of *Arabidopsis* grown under low Mg^2+^ were halved in comparison to controls (Fig. **3**). As a second proof of concept experiment, NaCl was added to the rooting medium. A continuous accumulation of apoplastic Na^+^ and Cl^-^ over the duration of NaCl exposure was observed (Fig. **4**). This is an anticipated consequence of salt stress, which has previously been reported as endpoint measurement only (Shahzad et al., 2013; Franzisky et al., 2021). The verification of these anticipated physiological responses confirmed the suitability of the new AWF extraction method for depicting mineral composition in the leaf apoplast over time.

The above mentioned NaCl-stress experiment was also used to test the new methods suitability to follow changes in the leaf apoplastic metabolite pattern over time. While there is plenty of knowledge on the effect of salt treatment on metabolite shifts in the symplast, information about leaf apoplastic metabolite dynamics are rare. In the leaf cells of salt-sensitive species, the metabolic acclimation to salt comprises changes in the levels of compatible solutes such as proline and other stress responsive metabolites such as other amino acids, sugars and small organic acids (Sanchez et al., 2008; Franzisky et al., 2021). However, salt stress-related changes in apoplastic metabolites have hardly been reported, except for a combined treatment by Na^+^ and alkaline pH (Sagervanshi et al., 2022), which led to increases of apoplastic sugars and small organic acids. The lack of information on changes of apoplastic metabolites in response to salt stress might be due to the difficulty in extracting minimally contaminated AWF from plants that might have leakier membranes as a result of salt exposure (Demidchik et al., 2014). Using the new AWF extraction method, minimally contaminated AWFs could be collected from *V. faba* after up to seven days of mild NaCl stress (Fig. **2**). The general pattern of the metabolic response to NaCl observed in the leaf apoplast of both species (Fig. **4**) resembled those previously observed in leaf tissue (Sanchez et al., 2008; Franzisky et al., 2021). This convergence between the apoplast and symplast suggests that the acclimation of the cellular metabolism translates into apoplast metabolite composition, thus highlighting the stark control of apoplast solutes by the symplast.

By the temporal resolved acquisition of leaf apoplastic metabolite, we demonstrated that following one day of NaCl exposure, beta-alanine or γ-aminobutyric acid (GABA) exhibit an increasing trend within the apoplast, and continue to accumulate steadily throughout the experimental period of NaCl exposure (Fig. **5**). Although the stress-responsive accumulation of GABA is a reaction observed in leaves of many plant species (Kinnersley and Turano, 2000; Che-Othman et al., 2020), to our knowledge, the salt-induced GABA accumulation in the leaf apoplast has not been demonstrated so far. In addition to the accumulation of various amino acids, two flavonol bisglycosides, most likely kaempferitrin (3,7-dirhamnoside kaempferol) and kaempferol 3-O-glucoside-7-O-rhamnoside, accumulated in response to the salt exposure in *V. faba*. Kaempferol glycosides are present in leaves of various plant species (Bozzo and Unterlander, 2021), including *Arabidopsis* (Unterlander et al., 2022) and *V. faba* (Weissenbock et al., 1984). Although flavonoid conjugates are found primarily in the symplast, e.g. the vacuole, chloroplast or nucleus (Zhao and Dixon, 2010), there is vague evidence for an occurrence in the plant cell wall as well. For example, kaempferol 3-O-glucoside in the apoplast of spruce and pine needles (Schnitzler et al., 1996; Strack et al., 1988), kaempferol bisglycosides in the leaf apoplast of *Arabidopsis* (Booker et al., 2012), and kaempferol glucosides derivates in the apoplast of lisianthus flower petals (Markham et al., 2000). Anticipated functions of flavones located in the epidermal cell wall are UV light protection, regulation of auxin transport and lignification of the cell wall (Jansen et al., 2001; Cesarino, 2019). However, the role of glycosylated flavones such as kaempferitrin in the cell wall is not yet well understood. By highlighting the current knowledge gap regarding the functions of flavonoids in the apoplast (Zhao, 2015; Bozzo and Unterlander, 2021), our new method offers a promising approach for assessing the role of apoplastic flavonoids.

### 4.4 Mapping hormones in leaf apoplast microdomains

To gain insights into stress signaling, levels of stress-responsive phytohormones such as ABA were investigated. The accumulation of ABA in leaves is known to occur time- and tissue-specific (Fricke, 2004). From this it can be postulated that ABA concentrations may differ between different leaf apoplastic domains. Yet, there are few methods that allow non-invasive, spatial mapping of plant hormones such as a Förster resonance energy transfer (FRET) biosensor but only for cytosolic measurement of ABA (Jones, 2016). However, using our new method it was possible to analyze the ABA concentration in the apoplast over several days in different apoplastic leaf domains to reveal spatiotemporal resolved ABA responses to NaCl exposure. After stressing *V. faba* for one day with NaCl treatment, ABA increased in the apoplast of the leaf base rather than the tip (Fig. **6**), suggesting that ABA dynamically accumulates in distinct parts of the leaf apoplast (Fricke, 2004). Such a microdomain-specific hormone response was shown for IAA, which decreased in the apoplast of the leaf tip rather than the base after one day of NaCl exposure (Fig. **6**). This means that in the field of plant biology it is worthwhile to explore the presence of spatial gradients of solutes within the leaf apoplast. Such leaf apoplastic ABA microdomains could be relevant for establishing growth zones and non-elongation zones within the same leaf (Fricke, 2004), a fact that is related to changes in the ABA concentration under salt stress (Fricke et al., 2004). Overall, the method presented overcomes various shortcomings of the ‘traditional’ methods to extract apoplastic solutes: it allows to analyze apoplastic solutes (i) minimal-destructively and (ii) over time. Moreover, (iii) apoplastic solute gradients between leaf microdomains can be examined. Thus, this approach is a valuable tool that enables the simultaneous capture of a wide range of compounds such as ions, metabolites, proteins, hormones and others with just a single sampling event. As a result, it advances the study of the role of the leaf apoplast, including its function as a refuge for microorganisms, nutrient store, hormone function, cell-cell communication or conduit for metabolite trafficking.

## 6. Author contributions

“J.K., M.S., C.X. and BL.F. conducted the experiments. BL.F., M.S., CM.G. and J.K. analyzed the data. CM.G. and K.H. designed the experiments. BL.F., CM.G., M.S. and K.H. wrote the paper.

## 7. Acknowledgments

We thank Lisa Mersmann, Jette Michalski and Elias Nehring for assistance in sample collection, Ralph Lehnart and Anja Giehl for technical support in ion measurement, and Dr. Joachim Kilian and Dr. Edda von Roepenack-Lahaye for extensive support in mass spectrometry.

## Supplementary figure captions

**Figure S1:**
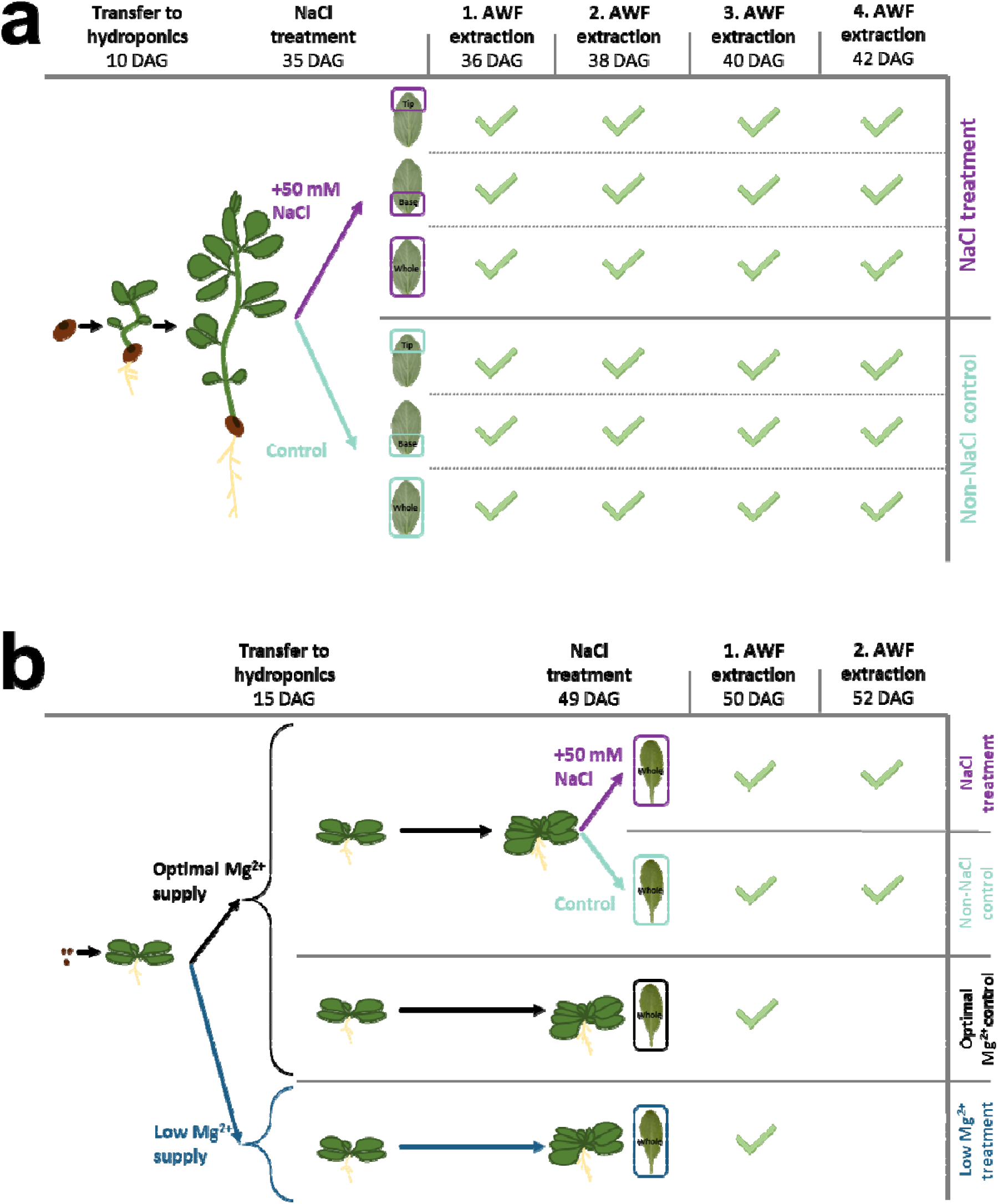
Overview of experiments performed with *Vicia faba* and *Arabidopsis thaliana*. A salt stress experiment was conducted with *Vicia faba* (**a**). For this, 30 mM NaCl was added to the nutrient solution, while the control received no NaCl. AWFs were collected from identical leaves on consecutive days, 35, 37, 39 and 42 days after germination, corresponding to the first, third, 5^th^ and 7^th^ days after the addition of NaCl. From controls and salt-treated plants, AWFs were collected from the whole leaf, i.e., the entire leaflet, as well as from the leaf tip or leaf base only (see Fig. 1d). Leaf tip and base are referred to as leaf microdomains. Two experiments were conducted with the *Arabidopsis* plants (**b**). First, *Arabidopsis* plants were grown with low magnesium (Mg^2+^) supply to demonstrate that the Mg^2+^ concentration in the leaf apoplast depends on the nutrients supplied at the roots. For this purpose, control *Arabidopsis* plants received optimal nutrient supply (0.5 mM Mg^2+^; see full recipe in section 2.1), while the treatment group received low Mg^2+^ supply (0.125 mM Mg^2+^). After 50 days since germination, AWFs were collected from whole leaves. Second, a group of *Arabidopsis* plants was exposed to salt stress by adding 50 mM NaCl to the nutrient solution. The salt stress experiment was done to demonstrate that the addition of NaCl to the rooting medium causes accumulation of Na^+^ and Cl^-^ in the leaf apoplast as a function of stress exposure, metabolic acclimation, and hormone responses over the course of the initial salt stress period. The control group remained without addition of NaCl. AWFs were collected from identical leaves on consecutive days, 50 and 52 days after germination, corresponding to the first and third days after the addition of NaCl. (DAG, days after germination)

**Figure S2:**
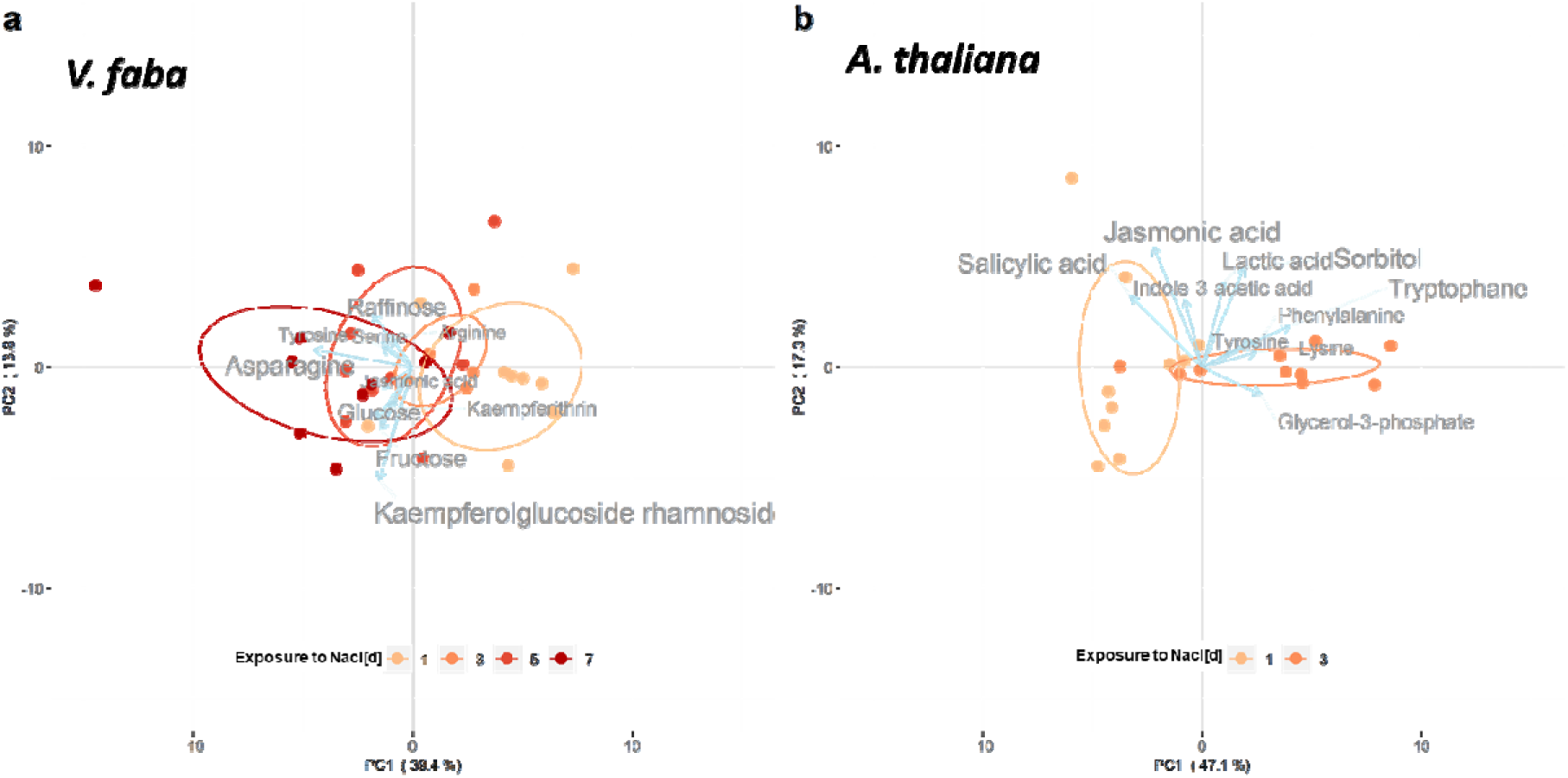
Unsupervised analysis of apoplast metabolite content in the apoplastic wash fluids (AWFs) of *Vicia faba* and *Arabidopsis thaliana* leaves in response to NaCl exposure over consecutive days. Principal component analysis (PCA) of the relative metabolite content in leaf AWFs of *V. faba* (**a**) and *A. thaliana* (**b**). Data were normalized over non-NaCl-stressed control (i.e. fold change_log2_(NaCl/Control)). Principal components (PCs) represent 53.2% and 64.4% of the total variance of the *V. faba* and *A. thaliana* data, respectively. PC 1 reflects the metabolic response to NaCl, i.e. the NaCl stress dose. Top 10 influential metabolites of PC loadings are indicated by labels and blue arrows. The segment length and the size of the metabolite labels correspond to the influence on the separation. Ellipses represent 0.5 confidence level (n = 6-8).

